# Normative sleep spindle database and findings from 772 healthy children from birth through 18 years

**DOI:** 10.1101/2022.03.31.486476

**Authors:** Hunki Kwon, Katherine G. Walsh, Erin D. Berja, Dara S. Manoach, Uri T. Eden, Mark A. Kramer, Catherine J. Chu

## Abstract

Work in the last two decades has identified sleep spindles, discrete “sigma band” oscillations during stage 2 sleep, as a key oscillatory mechanism required for off-line memory consolidation. Although, sleep spindles are known to evolve concomitant with brain maturation and reflect cognitive function across the lifespan, the details of this developmental trajectory are unknown. To address this, we curated a database of sleep electroencephalograms from 772 developmentally normal children to characterize spindles from birth through 18 years. After validating an automated spindle detector against ~20,000 hand-marked spindles across ages, we demonstrate that sleep spindle features follow distinct age-specific patterns in distribution, rate, duration, frequency, estimated refractory period, and inter-hemispheric spindle lag. These data expand our current knowledge of normal physiological brain development and provide a large normative database to detect deviations in sleep spindles to aid discovery, biomarker development, and diagnosis in pediatric neurodevelopmental disorders.

## 1. Introduction

Sleep spindles, discrete 0.5-2 second bursts of 9-15 Hz “sigma band” oscillations, are a hallmark feature of electroencephalogram (EEG) recordings during stage 2 non-rapid eye movement (N2) sleep ^1, 2^. Sleep spindles are generated by GABAergic neurons in the thalamic reticular nucleus and elaborated by well-delineated thalamocortical circuits ^3, 4^. Work in the last two decades has identified spindles as a key oscillatory mechanism required for off-line memory consolidation during N2 sleep ^5, 6, 7, 8, 9, 10, 11, 12^. Sleep spindles gate dendritic calcium shifts required for synaptic plasticity ^13^ and coordinate the reactivation patterns of sharp-wave ripples ^7, 8, 14^, hippocampal oscillations that reflect neuronal replay ^15^. Human cognitive studies have linked spindles to sleep-dependent consolidation of both procedural ^16, 17, 18, 19, 20, 21^ and declarative ^22, 23, 24, 25, 26^ memory tasks. Spindle rate correlates with general cognitive abilities and sleep dependent memory consolidation in both typically developing children ^27, 28, 29^ and those with neurologic disorders, including autism ^30, 31^, developmental delay ^30, 32^, and epilepsy ^33^. Confirming a mechanistic role, interventions that increase spindle activity result in improved sleep dependent memory consolidation ^34, 35, 36, 37, 38, 39, 40, 41^. Thus, sleep spindles provide a powerful non-invasive neurophysiological biomarker for thalamocortical circuitry and cognitive function across development.

Sleep spindles emerge early in life. In newborns, immature “pre-spindles” appear before 3 weeks of age^42^, followed by robust and prominent asynchronous sleep spindles between 3 and 9 weeks of age ^43^. These prominent spindle oscillations evolve dynamically in rate, duration, frequency, and inter-hemispheric synchrony, with the most marked changes observed in the first year of life ^32, 44, 45, 46^. Sleep spindles become increasingly synchronous over the first two years of life, reflecting the time course of cerebral white matter myelination ^32, 47, 48, 49^, and spindle frequencies are thought to increase linearly from childhood to adolescence in the frontal regions ^2, 50^.

Human spindles have historically been discriminated into two main types: slow spindles (<13 Hz) that predominate in frontal brain regions, and fast spindles (>13 Hz) that predominate in central and parietal brain regions ^45, 51, 52, 53, 54^. Previous studies have suggested that these different spindle populations have different cognitive roles, with frontal slow spindles more consistently related to declarative memory tasks^10, 26, 55^ and central fast spindles related to motor learning ^10, 56, 57, 58^. The reported frequency border between fast and slow spindles varies across ages and studies ^10, 45, 53, 59, 60, 61^, and when these distinct populations emerge over development is unknown.

While sleep spindles offer a promising biomarker of cognitive development and neurological disease across the lifespan, careful study of the developmental dynamics and normative values for spindle features are lacking for pediatric age groups. Previous work describing spindle features over development have been limited to clinical descriptions based on visual analysis or quantitative studies with small sample size ^45, 46, 62, 63^, limited electrode coverage ^50, 62, 63, 64^, low age resolution ^2, 64^, and limited pediatric age ranges ^2, 50, 53, 63, 65^. The challenge of obtaining sleep EEGs in healthy pediatric subjects and the lack of appropriate automated spindle detectors robust to age-specific spindle features impede development in this area.

To address this gap, we curated and analyzed a database of scalp EEG recordings during N2 sleep from a large cohort of developmentally normal children from age 0 to 18 years. We then trained and validated an automated spindle detector with hand-marked spindles across age groups. Using this detector, we evaluated several sleep spindle features—rate, duration, percentage, peak frequency, minimum inter-spindle interval, and interhemispheric synchrony—across typical development. We found sleep spindle characteristics follow distinct age-specific and regional-specific patterns. These data expand our current knowledge of normal physiological brain development and provide a large normative database to aid discovery, biomarker development, and diagnosis in brain development and pediatric neurodevelopmental disorders.

## 2. Materials and Methods

### 2.1. Participants and EEG recordings

Subjects aged 0-18 years with normal EEG recordings (as defined by clinical electroencephalographers independent from this study) in which N2 sleep was captured, were retrospectively identified from recordings performed in the Massachusetts General Hospital EEG laboratory between February 2002 and June 2021 (excluding July 2012-March 2015 due to data storage issues). Clinical chart review was performed and only those children with documented normal neurodevelopment and events leading to EEG evaluation that are not expected to alter EEG rhythms or cognitive function were included. Children that received neuroactive medications during the recording period were excluded. Children with chronic neurologic or psychiatric diagnoses were excluded, with the following exceptions: children diagnosed with mild attention or depression symptoms or tics not requiring medication treatment were included. Children with provoked seizures (e.g. not epilepsy, but seizures due to syncope, hypoglycemia, or other transient metabolic derangement), including febrile seizures, were included as the risk of subsequent epilepsy is similar to the general population ^66^, but we note we also evaluated this cohort separately. Children born prematurely (<37 weeks gestational age) were excluded. In addition, EEGs from 27 children ages 6-18 years recruited as population controls in research protocols at our institution were included. Of 819 total EEG recordings, 2 (0.2%) had poor recording quality and were excluded; 36 (4.4%) had < 3 min of N2 sleep and were excluded, resulting in 781 EEG recordings from 772 unique children for analysis.

Clinical EEG data were acquired following the international 10-20 system for electrode placement (Fp1, Fp2, F3, F4, C3, C4, P3, P4, F7, F8, T3, T4, T5, T6, O1, O2, Fz, Cz and Pz) with a standard clinical recording system (Xltek, a subsidiary of Natus Medical). Sampling frequency varied from 200 to 512 Hz. Research EEG data were acquired using a 64-channel cap (Easycap) and the 10-20 channels were selected for analysis. Research EEG data were acquired at 2035 Hz and down-sampled to 407 Hz. In each case, impedances were maintained below 10kΩ. All EEGs were visually reviewed by a board-certified pediatric neurophysiologist (CJC). Channels with poor recording quality and periods of significant artifact were ignored. Epochs containing N2 sleep (or trace alternant (TA) or quiet sleep (QS) patterns for <1 month old subjects) were identified according to standard criteria ^67, 68^. The N2 EEGs were then re-referenced to an average signal for subsequent analysis. All data analyses were conducted in accordance with protocols approved and monitored by the local Institutional Review Board according to National Institutes of Health guidelines.

### 2.2 Manual spindle detections

Two reviewers trained in spindle detection across ages performed manual spindle markings by consensus on 93 healthy subjects ages 0-18 years old. Following standard criteria, spindles were required to last a minimum of 0.5 seconds ^67, 68^. To train an automated spindle detector robust to both healthy and disease states for subsequent use, we also included manual spindle markings from separate projects including 10 subjects with continuous spike and wave sleep with encephalopathy ages 3-18 years old, and 12 subjects with Rolandic epilepsy ages 4-15 years old, in our training dataset ^33^. In each case, 100 s of EEG data from 19 channels using the standard 10-20 EEG montage were manually reviewed and the start and end time of each spindle marked, resulting in 19,625 total manually marked spindles from 115 unique pediatric subjects.

### 2.3. Sleep spindle detector

Typical N2 sleep architecture includes both vertex waves and K-complexes, brief events with increasingly sharper slopes at younger ages ^67^. Such sharp events typically present by age 5 months and produce wideband spectral features ^69^ that impact measures of sigma power common in many spindle detection approaches ^2, 33, 70^. For accurate detection of spindles in the setting of sharp sleep architecture, we adapted an automated latent state model (LS) spindle detector that we developed specifically to perform well in the setting of sharp events in the EEG ^33^. A detailed description of the detector can be found in 33 and code to apply the LS spindle detector is available for reuse and further development at https://github.com/Mark-Kramer/Spindle-Detector-Method. Application of this detector to this pediatric dataset consisted of three steps: 1) training; 2) validation across age groups; and 3) application. For each step, for each channel, we evaluate 0.5 s intervals of data and compute three EEG features: theta band power (4-8 Hz), sigma band power (9-15 Hz), and the Fano factor of the oscillation intervals—a measure of cycle regularity. We chose 0.5 s intervals, which are the typical minimum duration accepted for sleep spindles ^2, 12, 71, 72^, to maintain a 2 Hz frequency resolution, allowing reliable estimation of the theta band power. We advance each 0.5 s interval by 0.1 s, enabling detection of spindles at least 0.5 s in duration with 0.1 s resolution. To estimate the theta and sigma power in an interval, we detrended the (unfiltered) data, applied a Hanning taper, computed the Fourier transform multiplied by its complex conjugate, and divided the power at each frequency by the summed total power from 1-50 Hz. To compute the variability of oscillation cycles, we considered bandpass filtered data between 3-25 Hz (FIR, stop band attenuation 40 dB at 3 Hz, stop band attenuation 20 dB at 25 Hz, passband ripple 0.1 dB) and identified the peaks and troughs (minimum peak distance 28 ms, minimum peak prominence 2 uV); here filtering reduced the impact of high frequency activity on peak/trough detections. Then, to characterize the variability of the times between adjacent peaks and troughs we computed the Fano factor ^73^. We took the natural logarithm of each feature, shifted the interval by 0.1 s, and repeated these computations for the entire duration of the EEG signal for each channel.

To train the LS spindle detector, we used the manual spindle detections described in Section 2.2. For each channel (n=18 channels) and each subject (n=115), we computed for each 0.5 s interval the three features and assigned a spindle state label (“spindle” or “not spindle”). An interval was designated as “spindle” only if the entire interval lied within the bounds of a manually marked spindle. From these data, we fit empirical likelihood functions to each feature and state and estimated the transition matrix between the “spindle” and “not spindle” states. The LS spindle detector estimates the probability of a spindle in each 0.5 s interval, providing an easily interpretable value. Briefly, the probabilities of the spindle (p_yes_) and not spindle (p_no_) state are first initialized at 0.5. Then, for the first 0.5 s interval, the transition matrix is applied to [p_yes_, p_no_] to compute the one step prediction [p^1^_yes_, p^1^_no_]. The three features for this interval are then computed and the likelihood of each feature is used to compute the posterior [p^*^_yes_, p^*^_no_]. The posterior is normalized so that the probability sums to one (*i.e*., p^*^_yes_ + p^*^_no_ = 1), where p^*^_yes_ is the probability of a spindle for this interval. This process is repeated for each 0.5 s intervals in 0.1 s steps for the duration of the signal, where the [p_yes_, p_no_] for each interval is set to the normalized posterior of the previous interval.

To validate the LS detector across age groups, we performed a leave-one-out cross validation. To do so, we trained the detector with one subject omitted. We then applied this LS detector to estimate the probability of a spindle for each 0.5 s interval of each marked electrode of the omitted subject. We repeated this process for all 115 subjects. We then evaluated detector performance across all subjects using standard measures (see Statistical Analysis). We found that the probability threshold value of 0.95 is optimal across ages, so spindle detections were identified when probability of a spindle exceeded this threshold value (i.e., when the probability of a spindle exceeded 95%). Spindle detections separated by less than 1 s were concatenated ^2, 33^.

### 2.4. Analysis of spindle features

From the detected sleep spindles, the rate (mean number of spindles per minute), duration (mean duration of spindles in seconds), and percentage (percentage of time with spindles during the N2 recording) were calculated at each electrode for each EEG recording. To compute peak spindle frequency, we first estimated the power spectrum of each spindle. To do so, we applied a Hanning taper and 5 s of zero padding to each spindle detection and then estimated the power spectrum using the Fast Fourier Transform. At each spindle detection, the power spectra from 8-16 Hz were then divided by the summed power over 1-50 Hz. The resulting power spectra were averaged in each channel and each age group, and the peak frequency was identified as the frequency in the spectrum with the highest power (**Supplementary Figure 1**).

**Supplementary Figure 1.**
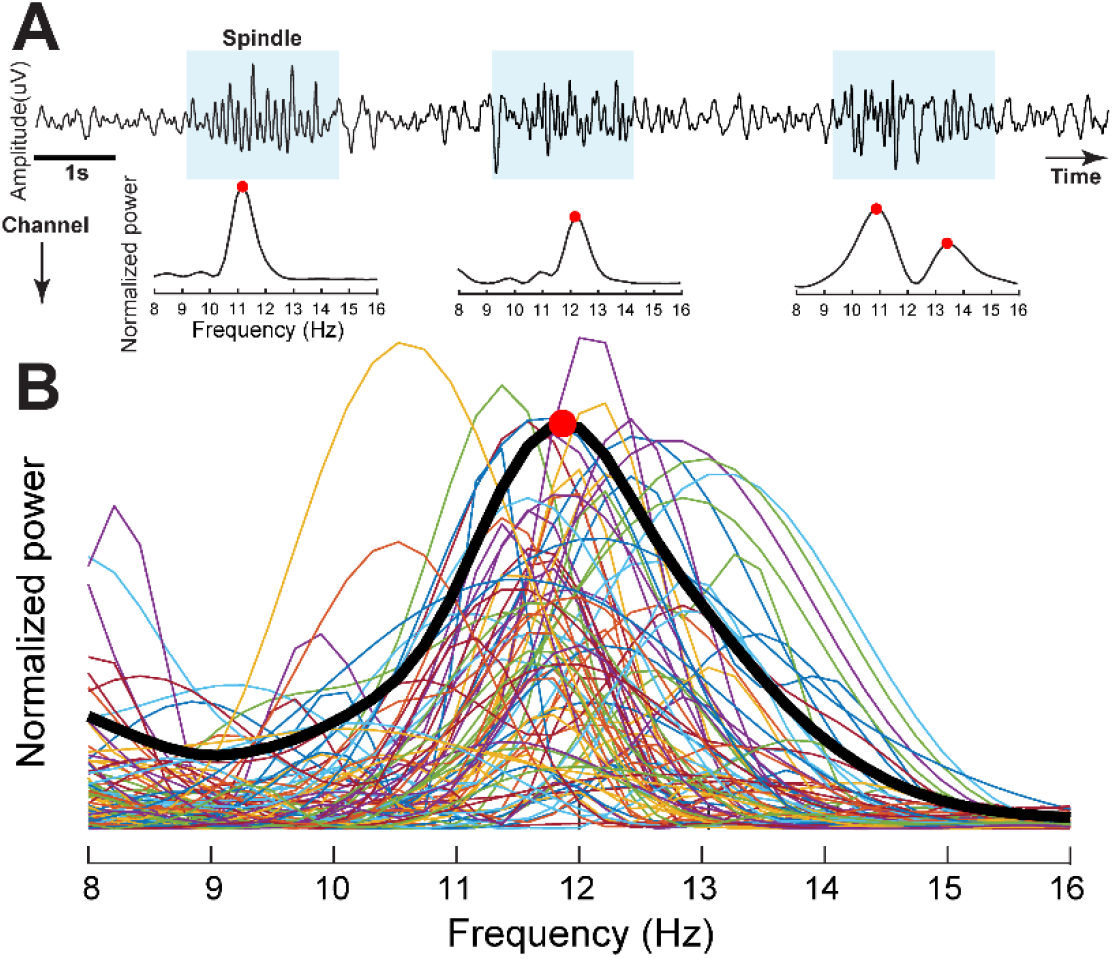
Spindle frequency analyses. **A)** An example 15 s recording from one channel showing spindle detections (blue). The normalized spindle spectra and peak frequencies (red markers) were identified. Note, some spindles demonstrate two peaks which will be retained in the distributions but not peak frequency analysis. **(B)** Example normalized spectra (thin colored curves) from electrode Fz from a 10 year-old subject. From the average spectrum (thick black curve) the peak spindle frequency (red marker) was identified.

The inter-spindle interval (ISI) was calculated as the time interval (s) between the end of a spindle and the onset of the next spindle in the same channel. We computed the ISI for the central electrodes from each EEG recording. To estimate spindle refractory periods, we computed the average of the lowest 10% of ISIs for each channel in which at least 10 spindles were detected. We choose the lowest 10% of ISIs to focus analysis on the shortest times between spindle detection while also minimizing the impact of outliers. To compute inter-hemispheric spindle lag, we first identified all times when spindles were detected in at least two frontopolar or centroparietal channels on either hemisphere (FP1, F3, C3, P3 or FP2, F4, C4, P4). The spindle lag was then calculated as the time between the first detected spindle among these channels in one hemisphere and the first detected spindle among the homologous channels in the opposite hemisphere.

### 2.5. Statistical analyses

To assess the performance of the spindle detector across ages, we report positive predictive value (PPV), sensitivity, and the F1 score (the harmonic mean of the PPV and sensitivity) of the detector relative to manual expert classification. To provide the most conservative assessment of detector performance, for each measure, we performed a by-sample analysis, in which manual and automated detections are compared at each sample of data ^33, 70^. For all subsequent measures, we used the detector threshold that optimized the F1 score (95%).

To characterize spindle features over age, subjects were categorized by year, rounding down to the most recent birthday. For visualization, topographic maps of spindle rate, duration, and percentage were interpolated using the Fieldtrip toolbox (http://www.ru.nl/neuroimaging/fieldtrip)^74^.

To model spindle features across age, the spindle features (except for minimum ISI) were first transformed using an inverse hyperbolic sine function. This transformation is a sigmoid function whose domain is the whole real line, which tends to reduce the magnitude of extreme values ^75^. Spindle rate, duration, and percentage did not follow a linear relationship with age. To characterize these features, we applied locally weighted scatterplot smoothing (LOESS) which uses weighted linear least squares and a quadratic polynomial model with a span of 0.3 ^76^. The 95% bootstrap pointwise confidence intervals of the LOESS fitted line were generated by resampling with replacement from the empirical data with 1000 iterations. The population 95% confidence intervals were generated for each age bin with a window size of 0.3, and values across immediately adjacent age bins were smoothed with a rolling average with a window size of 0.3. For minimum ISI and spindle lag, we applied a general linear model with the age as the predictor where p < 0.05 was considered as significant.

To test for differences in spindle rate between males and females at each electrode and each age group, we used a two-sampled t-test where p < 0.05 after Holm-Bonferonni correction ^77^ was considered significant.

We performed a cluster-based permutation test to identify clusters of electrodes with spindle features higher than the mean across electrodes for each age group. ^78, 79, 80^. To do so, for each subject in each age group, we a) subtracted the mean spindle value across channels from the value at each channel, resulting in positive and negative deviations. We then b) evaluated the distribution of these deviations across subjects to identify channels with higher spindle values compared to zero, using a one-tailed t-test. Channels below a critical alpha-level (p<0.05) were identified and joined in a cluster if they were adjacent based on the 10-20 electrode placement system. Next, c) a cluster-level statistic was computed as the sum of the absolute value of all t-values within a cluster. To determine which cluster-level statistics were unlikely to occur by chance, we used a cluster-based permutation test. Assuming spindle features are equally likely to occur above or below the mean, we randomly shuffled the sign of the deviations in each subject from a). We then performed steps b) and c) and computed the largest cluster-level statistic from the resampled data. This process was repeated 5000 times. Clusters from unpermuted data were considered significant if their cluster-level statistic exceeded the top 5% of the permutation distribution of 5000 samples (e.g., p<0.05). This process was performed for the spindle rate, duration, and percentage for each age subgroup.

To test for differences in spindle peak frequency between frontal and central electrodes at each age subgroup, we compared peak frequency values at Fz and Cz using a one-sampled t-test where p < 0.05 after Holm-Bonferonni correction was considered significant.

## 3. Results

### 3.1 Data characteristics

We analyzed 781 N2 EEG recordings from 772 unique subjects (382 F, 49.5%) ranging in age from 0 days to 18 years. Clinical characteristics are provided in **Table 1**. The final diagnosis for the events that led to the EEG evaluations are listed in **Table 2**. The average duration of N2 sleep (TA or QS) per EEG recording was 14.04 min (IQR 8.40-18.07 min).

**Table 1.**
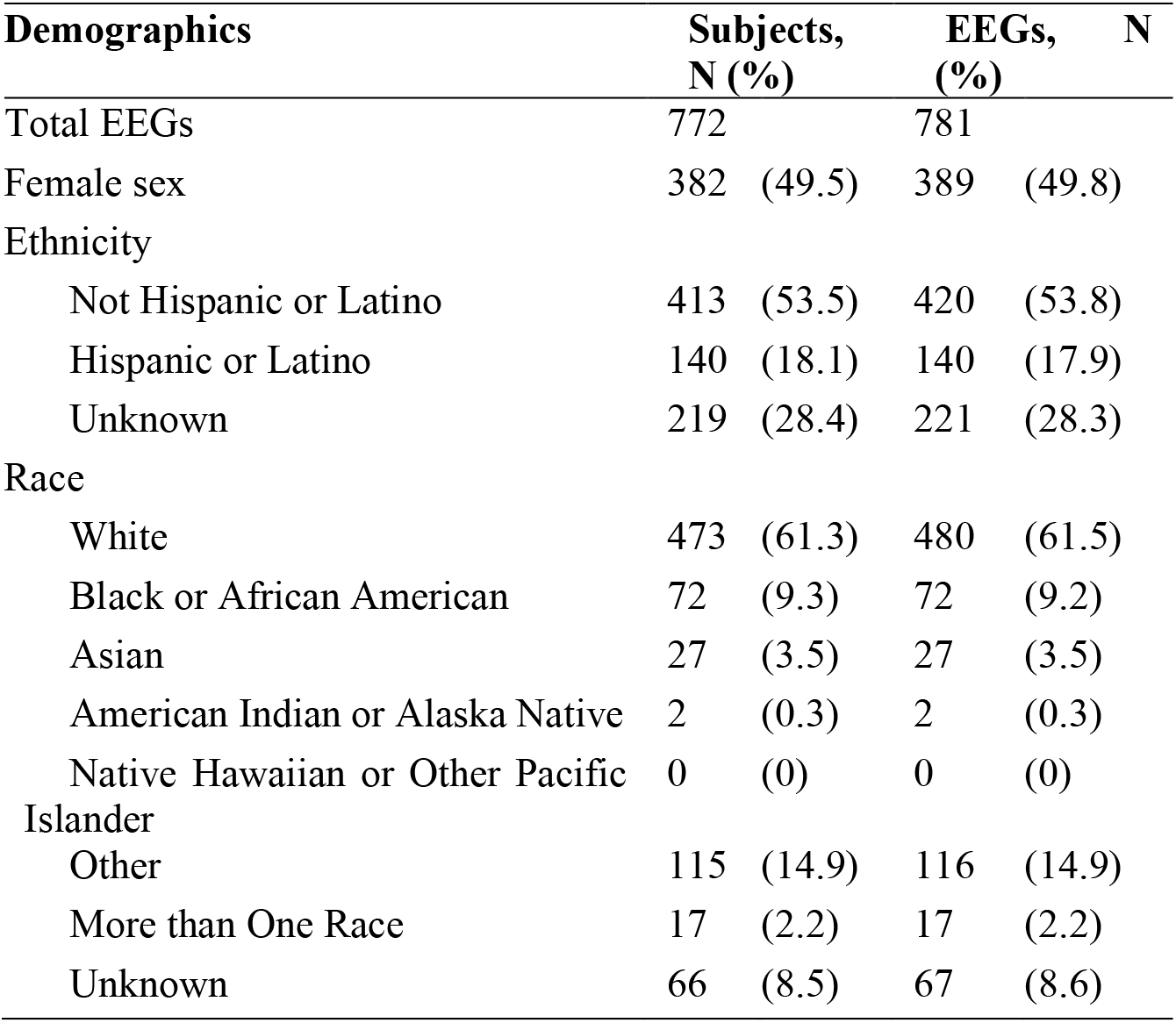
Patient demographics

**Table 2.** Indications for EEGs (N = 781)

| Diagnosis | N | Diagnosis | N | Diagnosis | N |
| --- | --- | --- | --- | --- | --- |
| Provoked seizure | 132 | FND | 15 | Mild hypertonia | 2 |
| Syncope/pre-syncope | 99 | Respiratory event | 14 | Muscle fasciculation | 2 |
| Migraine/headache | 57 | Tremors | 8 | Pain | 2 |
| Sleep phenomenon | 57 | Anxiety/panic attack | 7 | Aplastic anemia | 1 |
| Nonspecific movement | 56 | Dizziness | 7 | Bell's palsy | 1 |
| Staring spell | 44 | Altered mental status | 6 | Benign hypotonia | 1 |
| Gastrointestinal reflux | 30 | Concussion | 6 | Episode of unsteadiness | 1 |
| Stereotypy/tic | 30 | Vomiting | 6 | Hiccups | 1 |
| Behavioral event | 28 | Fatigue | 5 | Intussusception | 1 |
| Research control subjects | 26 | Toxic exposure | 5 | Lyme disease | 1 |
| Breath-holding spell | 23 | Vertigo | 5 | Polydipsia | 1 |
| Transient unresponsiveness | 23 | Visual phenomenon | 5 | Rigors | 1 |
| Unusual eye movement | 22 | Fall | 4 | Suspected abuse | 1 |
| Shuddering spell | 20 | Startle reflex | 3 | Transient weakness | 1 |
| BRUE | 18 | Hypoglycemia | 2 | Trauma | 1 |

The sleep phenomenon category includes all sleep related indications: sleep myoclonus (36), parasomnias (13), hypersomnia (1), periodic limb movement (1), somnolence (1) and other sleep phenomena (5). The provoked seizure category includes febrile seizures (108) and provoked seizures (24). BRUE: Brief resolved unexplained event; FND: functional neurologic disorder

### 3.2 Automated spindle detector validation across age groups

Leave-one-out cross validation of the automated spindle detector revealed excellent performance against manual markings in 93 normally developing children ages 0-18 years (n=47 infants ages 0-2 years old, n=46 school age subjects ages 2-18 years; **Figure 1**). Balanced detector performance was achieved using a 95% probability threshold (mean F1: 0.44, 95% CI [0.11 0.76]). At this threshold, the detector’s sensitivity was 0.48 and positive predictive value 0.47 against hand markings. We note that this performance is higher than previously reported spindle detectors, which have ranged from F1 0.2-0.43 ^70^. Example spindle detections across ages are shown in **Figure 1C**.

**Figure 1.**
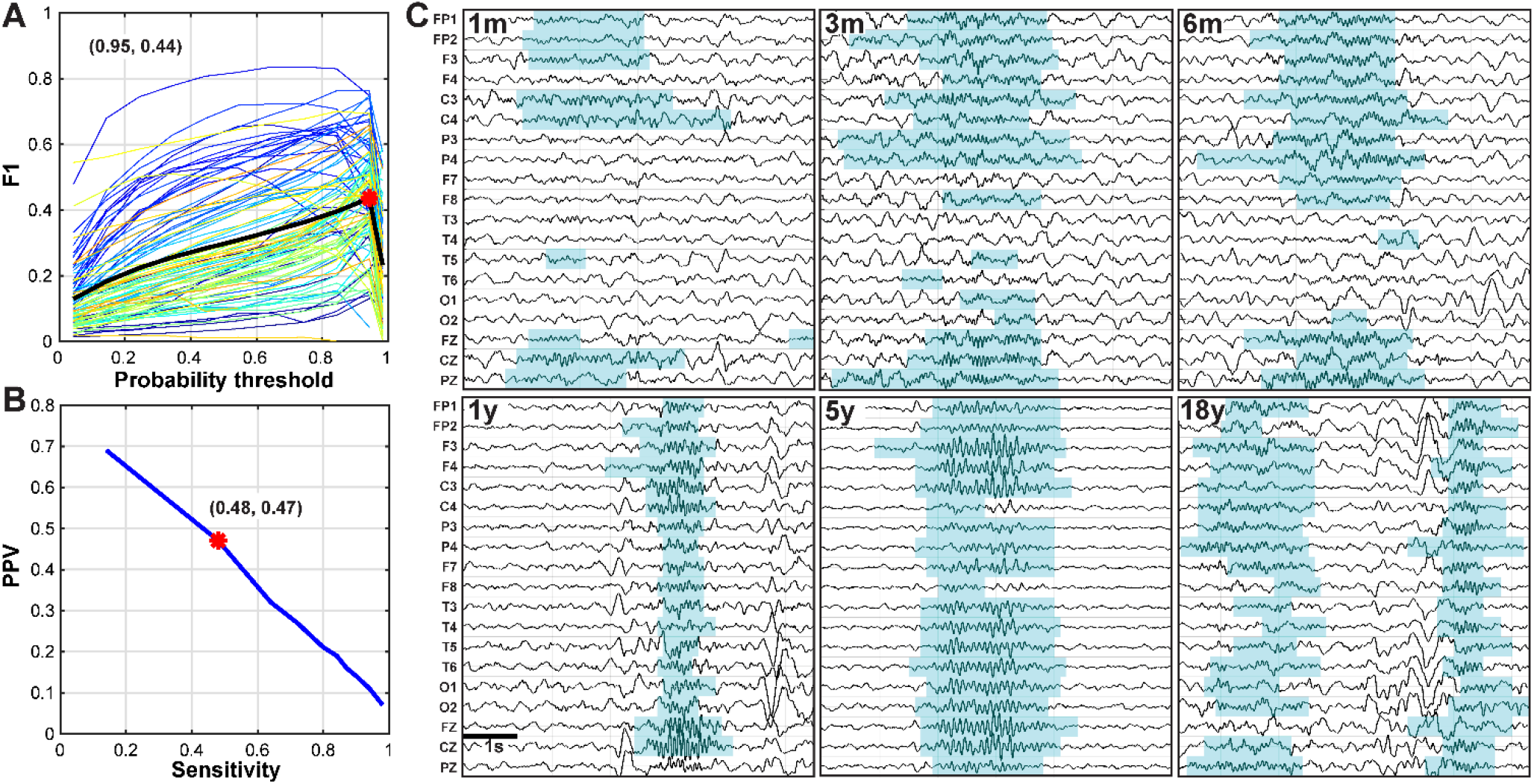
Spindle detector performance across ages. **A)** Using leave-one out cross validation, the optimal F1 statistic against hand markings was achieved at the 95% probability threshold (0.44, red marker). Colored curves represent detector performance for individual subjects across probability thresholds and the black curve indicates the mean performance across subjects. **B)** The sensitivity and positive predictive value (PPV) of the detector at different probability thresholds (95% threshold indicated in red). **C)** Example spindle detections in different age groups: 1 month, 3 months, 6 months, 1 year, 5 years, 18 years. Each sample shows five-seconds of N2 sleep with automated spindle detections highlighted in blue.

### 3.3. Evolution of normative spindle parameters over development

#### Spindle distribution

Sleep spindle characteristics follow a distinct, age-specific developmental trajectory. In infants, spindles are more prominent over frontopolar and central regions. In toddlers (1-2 years), spindles become more prominent over the central regions. Spindles then migrate anteriorly to become dominant in frontal regions by 5-6 years and frontopolar regions by 9-10 years, though with increasing rates appearing diffusely. Over adolescence, spindles migrate posteriorly and dominate in central and parietal regions by late teens (17-18 years; **Figure 2**).

**Figure 2.**
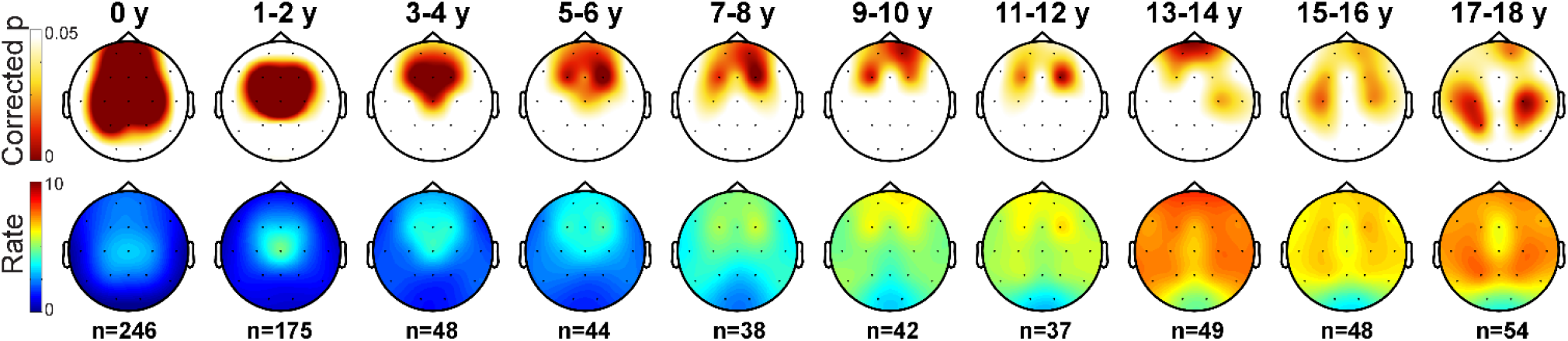
Topographic maps of spindle distribution over development. **Top row)** Electrode clusters with spindle rates higher than average (p<0.05) are shown. **Bottom row)** The mean spindle rate (number per minute) is plotted per age group. The number of EEG recordings included in each age group is indicated below each topoplot.

#### Spindle rate

Averaged across all channels, the mean spindle rate follows an N-shaped curve over development, with the most prominent nadirs at 0 months (0.47 spindles/min, 95% CI [0.25, 0.71]) and 2.34 years (1.40 spindles/min, 95% CI [1.17, 1.65]), and the most prominent peaks at 5.0 months (2.24 spindles/min, 95% CI [2.00, 2.51]) and 17.33 years (6.15 spindles/min, 95% CI [5.65, 6.70]). A secondary relative nadir is observed at 0.85 years (1.71 spindles/min, 95%CI [1.51, 1.93] and a secondary peak at 15 months (1.83 spindles/min, 95%CI [1.59, 2.11]). Spindle rate in the frontopolar (FP1/FP2), central (C3/C4), and parietal (Pz, P3/P4) regions follow this N-shaped developmental trajectory robustly, while spindle rate in the frontal (F3/F4, F7/F8) and anterior midline (Fz, Cz) regions follow a similar but less prominent trend. In contrast, spindle rate increases near-monotonically with age in the temporal (T3/T4), posterior temporal (T5/T6), with only subtle peaks observed at 0.42 and 1.56 years (0.73 spindles/min, 95%CI [0.56, 0.92]; 0.91 spindles/min, 95%CI [0.72, 1.12] in the temporal region, at 0.42 and 1.44 years (0.52 spindles/min, 95%CI [0.38, 0.66]; 0.98 spindles/min, 95% CI [0/78, 1.20] in the posterior temporal. In the occipital region, there is an initial nadir at 0.41 years (0.22 spindles/min, 95% CI [0.14, 0.31]) followed by a local peak at 1.39 years (0.72 spindles/min, 95% CI [0.56, 0.88]) (**Figure 3**).

**Figure 3.**
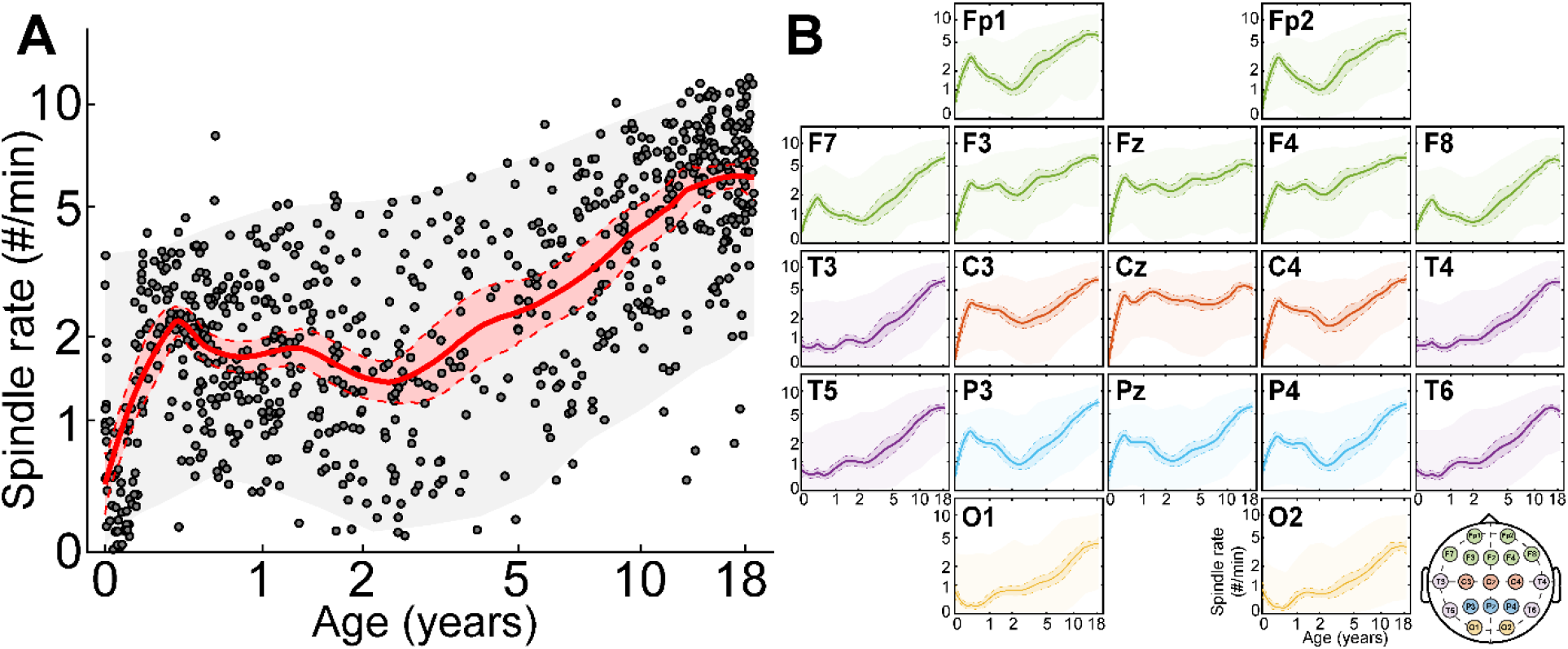
Normative spindle rates over development. **A)** Average spindle rate across all electrodes with age. The red line (shaded areas) indicates local regression fits (95% confidence intervals). The grey shaded areas indicate 95% confidence intervals of the population data. **B)** Results for each electrode, arranged according to 10-20 electrode placements. Colors indicate brain region (frontal-green, central (e.g., Rolandic)-red, temporal-purple, parietal-blue, occipital-yellow).

**Supplementary Figure 2.**
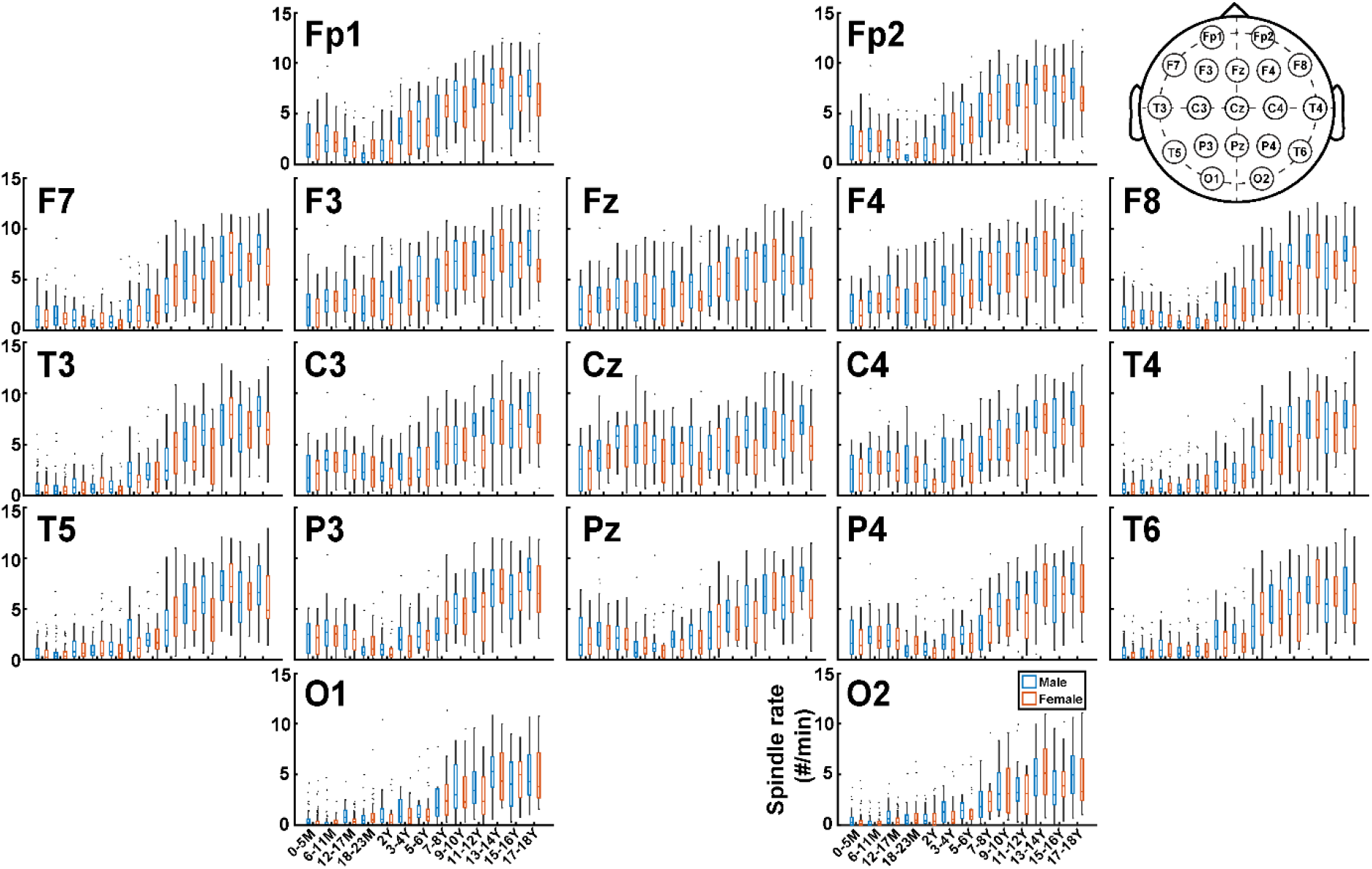
Spindle rate by sex across age and brain regions reveals no difference. Boxplots of spindle rate for male (blue) and female (red) subjects reveals no evidence of a difference in any brain region for any age group.

After correction for multiple comparisons, we found no evidence of a difference between male and female spindle rates in any region at any age (p>0.16 for all comparisons, **Supplementary Figure 2**). We note that children with a history of febrile seizures followed the same developmental trajectory as those without provoked seizures, and excluding these children produced equivalent results (**Supplementary Figure 3**).

**Supplementary Figure 3.**
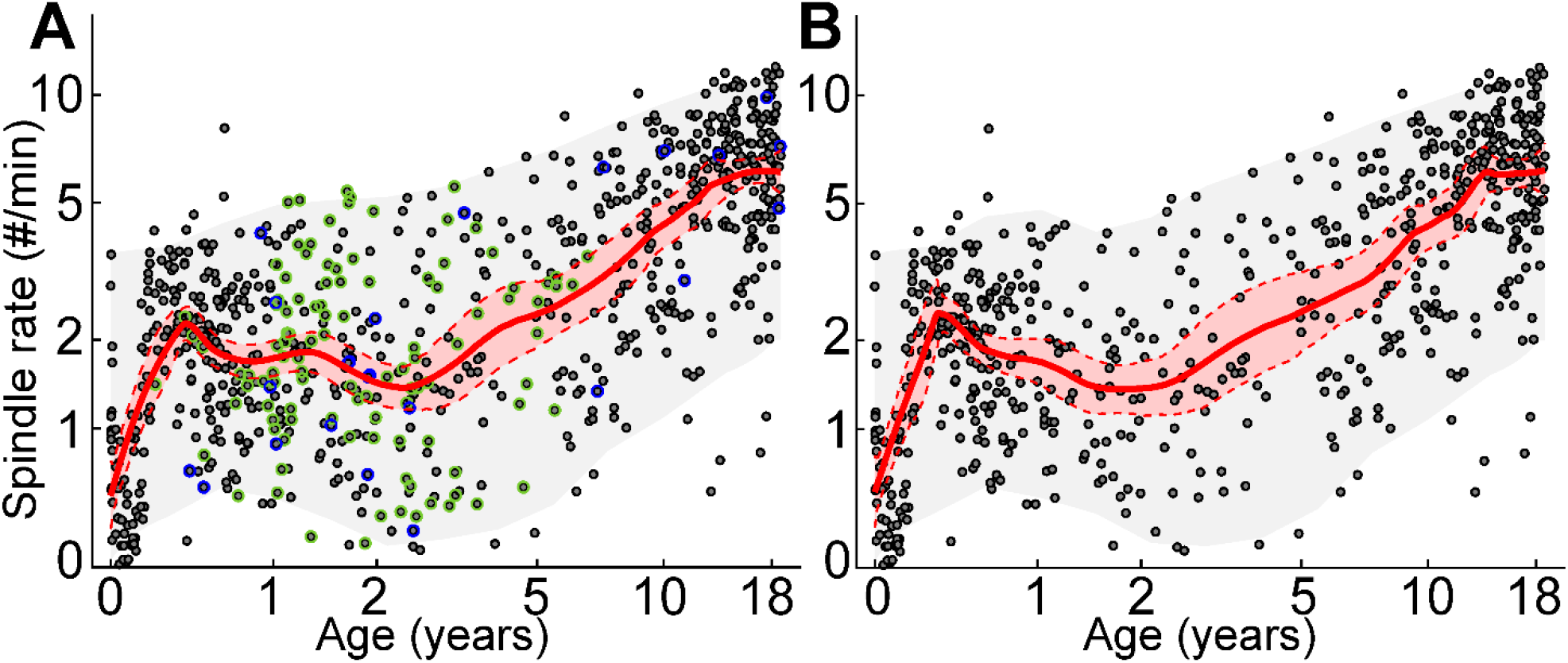
Children with provoked seizures have typical spindle rates. **A)** Spindle rate across all electrodes with age. Green (blue) circles indicate subjects with history of febrile (provoked) seizure; Grey dots are remaining healthy subjects. The red line (shaded areas) indicates local regression fits (95% confidence intervals). The grey shaded areas indicate 95% confidence intervals of the population data. **B)** Spindle rate over age after excluding subjects with febrile or provoked seizures follows the same trajectory as A).

#### Spindle duration

Changes is spindle duration follow a developmental trajectory similar to spindle rate. Averaged across channels, the longest spindles are present in mid-infancy at 0.39 year (mean 1.50 s, 95% CI [1.41, 1.59]) and late adolescence at 18 years (1.55 s, 95% CI [1.43, 1.67]). Nadirs are observed at 0 months (0.72 s, 95% CI [0.64, 0.80]) and 1.96 years (0.95 s, 95% CI [0.91, 0.99]). Like spindle rate, this N-shaped trajectory is again most prominent in frontopolar, central and parietal regions. In contrast, spindle duration increases consistently with age in temporal and occipital regions (**Supplementary Figure 4**).

**Supplementary Figure 4.**
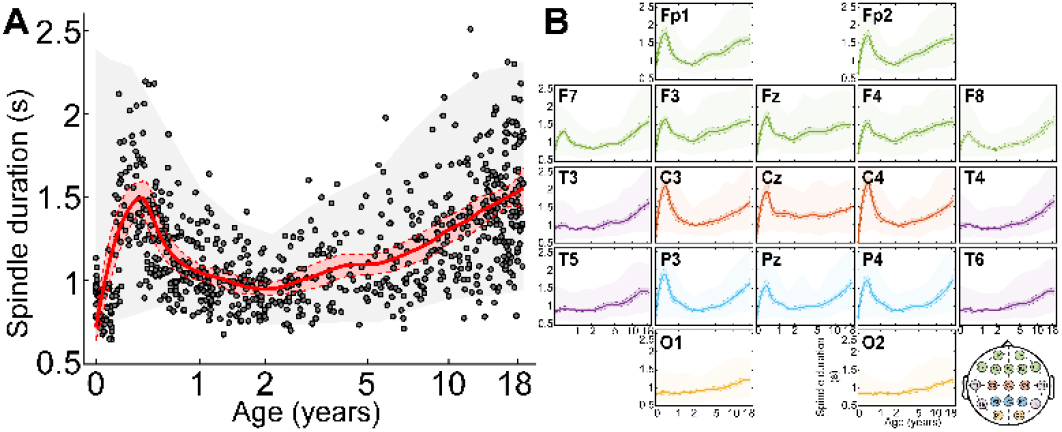
Normative spindle duration over development. **A)** Average spindle duration with age. The thick line indicates the local regression fit. The thick shaded areas indicate the 95% confidence intervals of the model. The lightly shaded areas indicate 95% confidence intervals of the population data. **B)** Similar plots are provided for each electrode, organized by the 10-20 electrode placement system. Colors are organized by brain region in the inset.

#### Spindle percentage

Spindle percentage (the percentage of time during N2 sleep with spindles) is an aggregate measure reflecting both spindle rate and duration. As such, the mean percentage of spindles across all channels follows a robust N-shaped curve over development, which is most prominent in central and parietal regions (**Supplementary Figure 5**). Averaged across channels, spindle percentage achieve local maxima at mid-infancy (0.41 year, mean 6.26%, 95% CI [5.20, 7.54]) and late adolescence (18 years mean 15.51%, 95% CI [12.53, 19.19]). A smaller peak occurs at 14 months (3.29%, 95% CI [2.79, 3.86]. Prominent nadirs were observed at 0 months (0.57%, 95% CI [0.23, 0.96]) and 2.30 years (2.37%, 95% CI [1.95, 2.87].

**Supplementary Figure 5.**
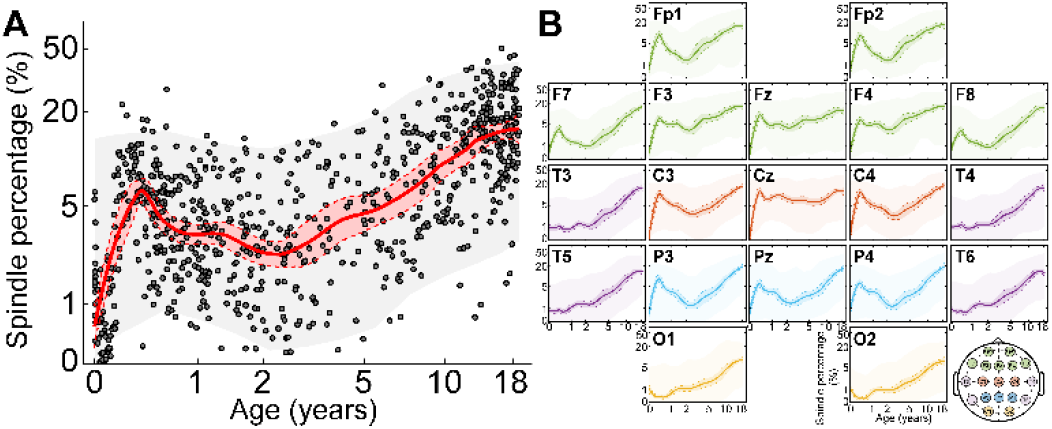
Normative spindle percentage over development. **A)** Average spindle percentage with age. The thick line indicates the local regression fit. The thick shaded areas indicate the 95% confidence intervals of the model. The lightly shaded areas indicate 95% confidence intervals of the population data. **B)** Similar plots are provided for each electrode, organized by the 10-20 electrode placement system. Colors are organized by brain region in the inset.

### 3.4. Spindle frequency over development

#### Spindle frequency changes with age

The peak frequency of spindles is highest in infancy and mid-adolescence, and similar between these age groups (0-1 year, mean peak frequency: 13.1 Hz; 14-18 years, mean peak frequency: 13.1 Hz). Between ages 1 and 2 years, there is a sharp drop in spindle frequency (2 years, mean peak frequency 11.2 Hz). From ages 2-14 years there is a consistent increase in spindle frequency with age. This U-shaped pattern of peak spindle frequency with age is present across all brain regions (**Figure 4)**.

**Figure 4.**
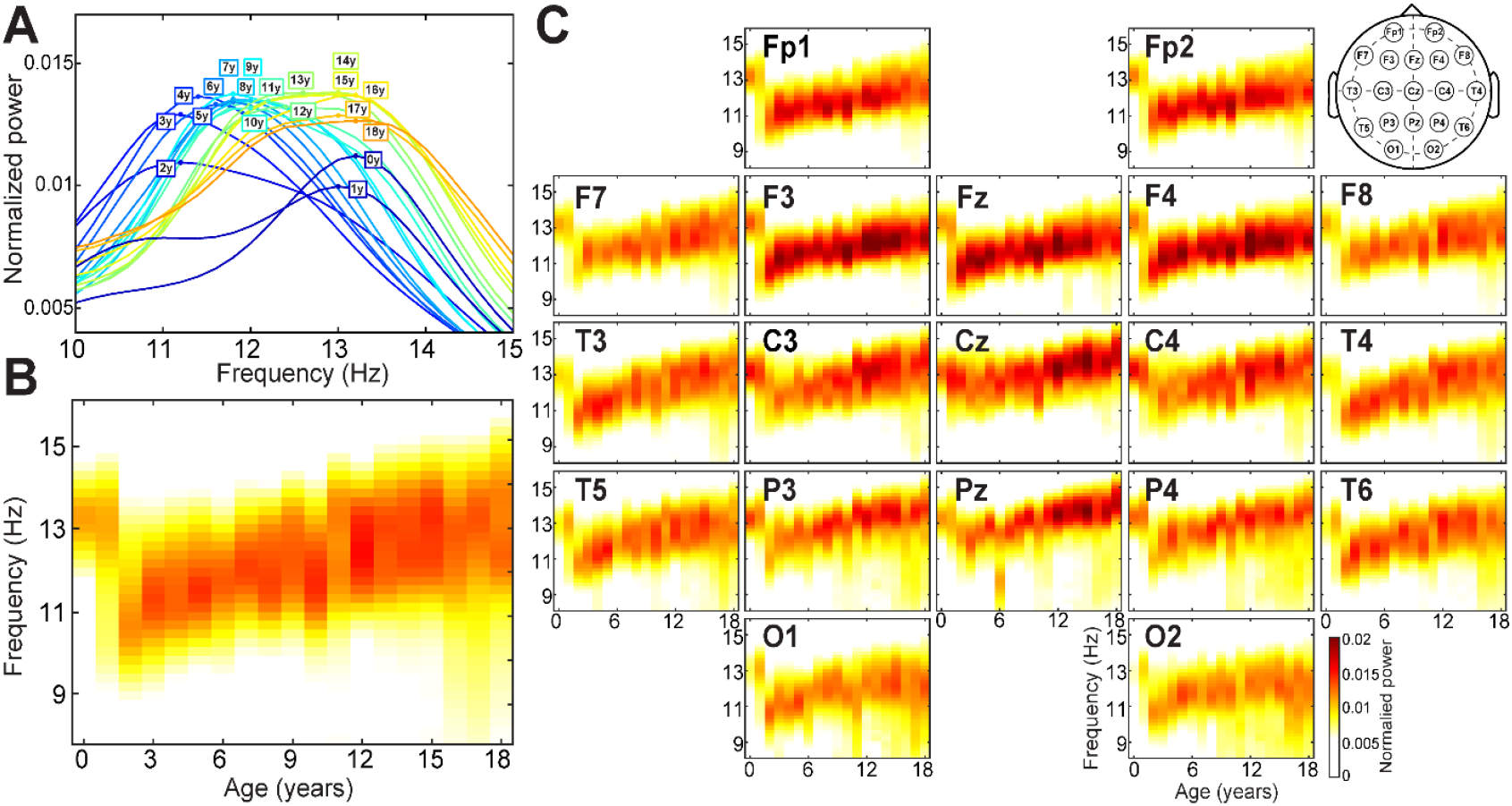
Normative spindle frequency over development. **A)** Each line represents the normalized peak spindle power across channels by age. Peak frequencies are high in infants and late adolescents. **B)** The distribution of normalized spindle frequencies across channels is plotted as a function of age. **C)** Spindle frequency distributions by age are provided for each 10-20 channel.

#### Slow and fast spindles emerge early in development

To identify when the canonical frontal “slow” and central “fast” spindles emerge and empirically characterize their frequencies, we compared peak spindle frequency between central (Cz) and frontal (Fz) regions over age. We found no difference between central and frontal spindle peak frequencies in patients <18 months (p > 0.23 for all comparisons). Beginning at 6 months, the peak frequency of frontal spindles declines and by 18 months frontal spindles had a lower peak frequency (11.7 Hz) compared to central regions (12.7 Hz; p<0.0001). After 2 years, peak frequencies in the frontal and central regions both increase with age through age 18 years, but remain distinct from each other, with lower frequencies always observed in frontal regions compared to central regions (**Figure 5,** p<0.05 for all comparisons after correction for multiple comparisons).

**Figure 5.**
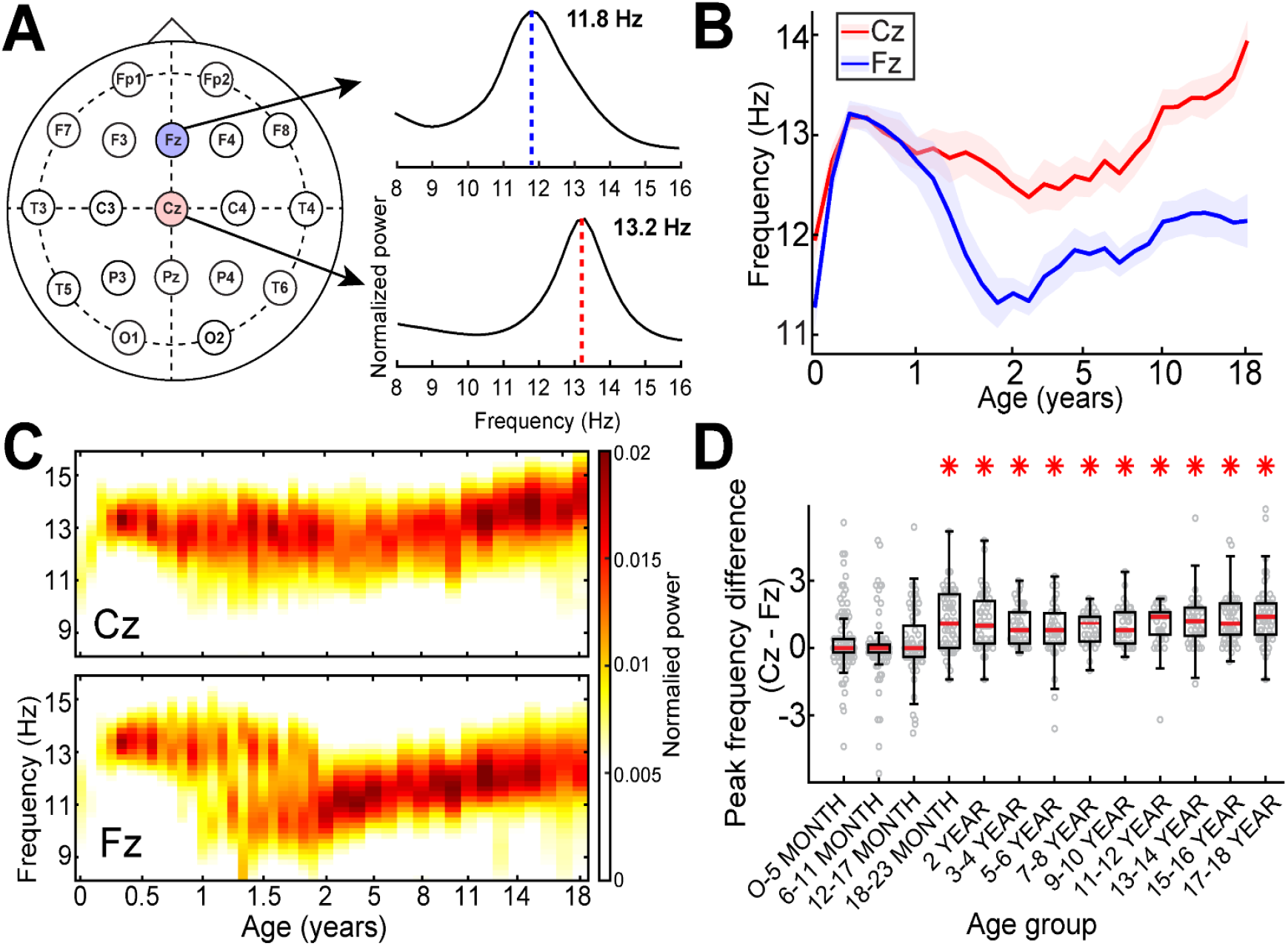
Frontal slow spindles emerge by 18 months. **A)** Example of averaged power spectra for spindle detections in Fz (blue) and Cz (red) in a 10 year-old. **B)** Peak spindle frequency in Fz (blue) and Cz (red) over 0 to 18 years. **C)** The distribution of normalized spindle frequencies across channels is plotted as a function of age. **D)** Spindle peak frequencies are lower in Fz compared to Cz channels after 18 months (* indicate p < 0.05 after correction for multiple comparisons).

### 3.5. Spindle refractory periods over development

To estimate spindle refractory periods over development, we computed the mean of the smallest 10% of inter-spindle intervals in the central regions in each hemisphere (channels C3 and C4). We found that the spindle refractory period decreases monotonically with increasing age (p=0, Beta= −1.15, 95% CI [−1.29, −1.02]), **Figure 6**).

**Figure 6.**
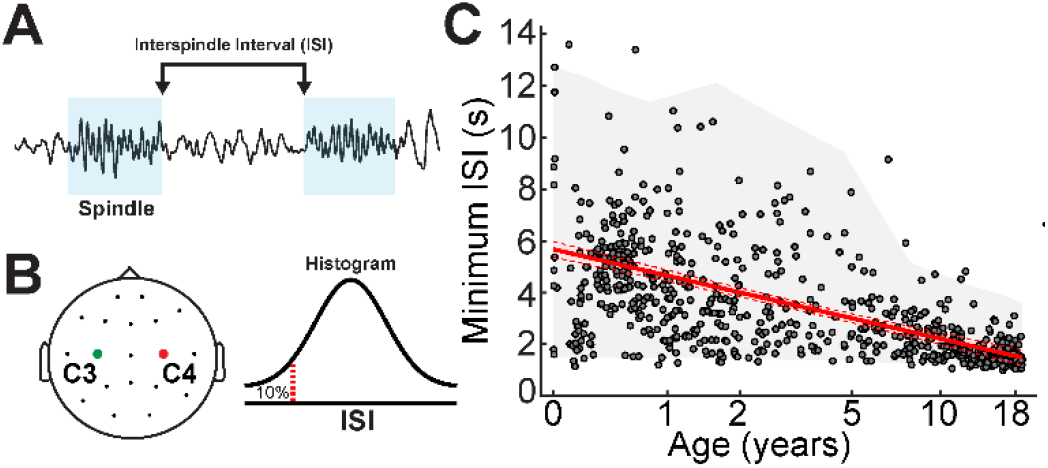
Minimum inter-spindle interval decreases with age. **A)** The ISI is calculated as the time from the end of one spindle to the start of the next. B) The lowest 10% of ISIs from channels C3 and C4 were considered. C). ISI decreases with age. The red line (shaded areas) indicates local regression fits (95% confidence intervals). The grey shaded areas indicate 95% confidence intervals of the population data.

### 3.6. Inter-hemispheric spindle lag over development

To characterize the coordination of spindles between hemispheres, we computed the inter-hemispheric lag between spindles detected in the frontal, central and parietal regions (channels FP1, F3, C3, P3 and FP2, F4, C4, P4; Figure 8A). We found that interhemispheric spindle lag decreases monotonically with increasing age (p=0, Beta = −0.28, 95% CI [−0.31, −0.24]) (**Figure 7**).

**Figure 7.**
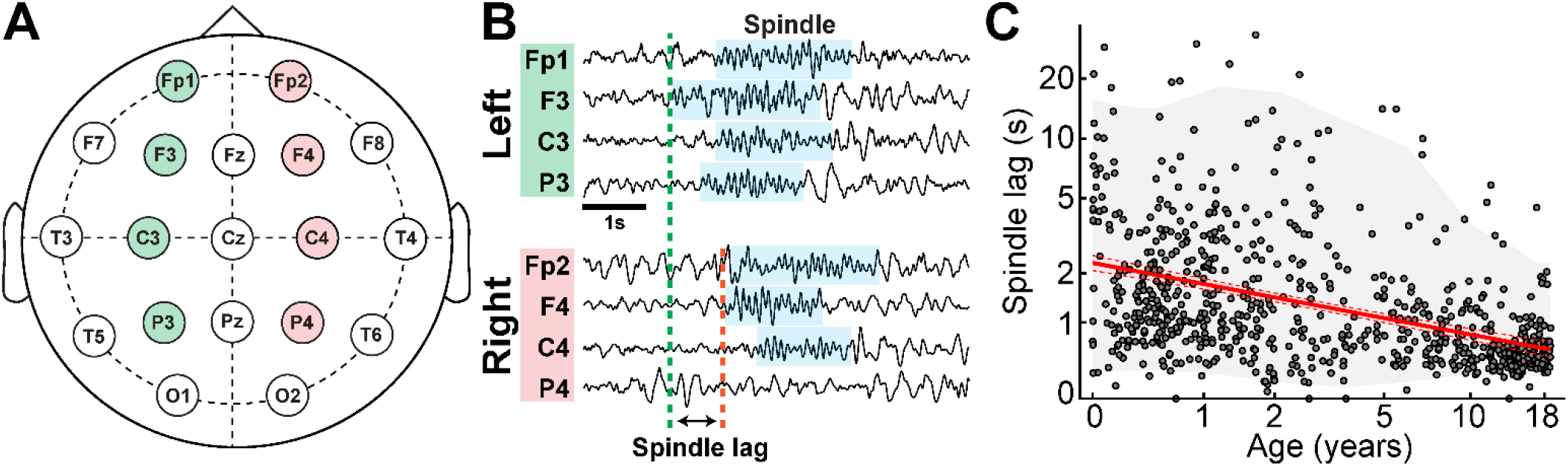
Inter-hemispheric spindle lag over development. **A)** Spindles detected in at least two frontocentral channels in the same hemisphere were analyzed. **B)** The spindle lag was calculated from the start of the first detected spindle in one hemisphere to the start of the first detected spindle in the contralateral hemisphere. **C)** Spindle lag decreases with age. The red line (shaded areas) indicates local regression fits (95% confidence intervals). The grey shaded areas indicate 95% confidence intervals of the population data.

## 4. Discussion

Spindles are fundamental thalamocortical sleep rhythms that are mechanistically linked to essential sleep-dependent cognitive processes. Although, sleep spindles are known to evolve concomitant with brain maturation, the details of this developmental trajectory have been lacking. By applying a validated automated spindle detector to a large database of N2 sleep EEGs from healthy neonates, infants, and children we characterize several key features of spindles over the course of development. This work reveals that, like anatomical brain maturation, sleep spindle features—including rate, duration, topography, refractory period, and inter-hemispheric synchrony follow a predictable, heterochronic, and non-linear developmental trajectory from birth to late adolescence. This novel neurophysiological data provides a direct window into the maturing thalamocortical networks that generate and drive these critical oscillations and insights into the development of the focal and distributed cognitive functions that they support.

The ability to utilize biomarkers to detect disease in pediatric populations is hampered by the dynamic changes in baseline measures that occur over healthy development. Using this large database, we provide robust normative parameters of spindle features for each age across development. Such high-resolution developmental natural history data can be used to assess for deviations from this distribution in at-risk populations to aid in diagnosis, elucidate thalamocortical pathophysiology, and measure treatment responses after interventions.

Using the same detection and recording techniques across ages, we found that very early “pre-spindles”^67, 81^, spindles observed in neonatal periods, and spindles observed over the first year of life share remarkably similar features to the mature spindles observed in late adolescents. Consistent with previous clinical observations in infants and a larger study of older children, we found that both spindles in infancy and late adolescence are dominated by faster ~13 Hz frequencies over central or Rolandic brain regions ^2, 67, 68^. In contrast, but also consistent with earlier work, we found that between infancy and adolescence, spindles are slower and dominate over frontal regions ^2^. As spindle frequency is a heritable trait ^2^, spindle measures from infancy may provide a reliable early window into the integrity of the thalamocortical circuitry required for mature spindles.

Compared to spindle frequency and topography, spindle rate and duration follow a different developmental trajectory, with an initial peak identified here and in previous smaller studies in the first year ^46, 47^ followed by a rapid decline and then steady increase from age two years onward ^2, 50^. Notably, these early non-linear dynamics rise and fall in striking parallel with the trajectory of cortical growth. Cortical grey matter increases dramatically (more than 100%) in the first year, due to ongoing neurogenesis, synaptogenesis, and increased spine density ^82, 83, 84^. Following this period of explosive growth, cortical thickness reverses course and begins to decrease between the first and second year, reflecting synaptic pruning ^85^. This U-shaped trajectory of cortical growth is heterochronic, with the most dramatic early cortical growth is seen in the central and parietal regions, the same locations we observe the most prominent spindle rates in the first year. In the prefrontal regions, cortical growth peaks around 18 months, which coincides with our detection of the first frontal slow spindles ^82, 86^. In contrast to gray matter, white matter maturation increases steadily over the first two years of life, then continues at slower rate through late adolescence ^85^. Thus, the near-linear change in spindle rate with age in older children observed here and in prior studies ^50, 54, 87^ parallels, and may reflect, ongoing maturation of thalamocortical white matter connections.

Spindle rate and duration are dictated by the thalamocortical circuits that generate them. In the setting of reduced noradrenergic and serotonergic signaling during N2 sleep, the relatively hyperpolarized GABAergic neurons in the thalamic reticular nucleus (TRN) can autonomously generate burst firing or can be triggered by relatively small glutamatergic inputs from thalamocortical neurons. The burst firing of inhibitory TRN cells results in hyperpolarization of thalamocortical neurons, which conversely activates a hyperpolarization-activated cation current (Ih) and disinhibits the T-type calcium current, leading to a burst of action potentials from the glutamatergic thalamocortical cells that target the cortex and sends feedback to further engage the TRN ^88^. How and why spindles terminate is thought to be related to after-depolarization refractory periods in the thalamocortical neurons. In *ex vivo* slices, spindles can persist indefinitely if the Ih current is blocked ^89, 90^. Interestingly, we observed the longest spindle durations during infancy, suggesting reflect an immature state of this current. We also observed the longest spindle refractory periods in infancy, suggesting that spindle duration and refractory periods are not simply anti-correlated and other factors must be contributing.

Several lines of evidence suggest that sleep spindles support off-line replay of learned experiences during sleep, resulting in improved consolidation of these memories ^9, 32, 87, 91, 92^. Several studies have linked sleep spindle rate with cognitive functions in children ^28, 29, 33, 93, 94, 95^ and adults in both health ^96, 97, 98, 99^ and disease ^30, 33, 61, 100, 101, 102, 103, 104^, with fast spindles more tied to procedural or motor learning ^56, 57, 58^ and slow spindles to declarative memory ^26, 55^. We identified that fast spindles are present from birth and distinct slow spindles consistently emerge by 18 months. This finding is consistent with clinical descriptions ^67, 105^ and our previous observations of discrete 13 Hz and 11 Hz bands emerging at ~16 months of age during non-rapid eye movement sleep (see Figure 2 in Chu et al, 2014 ^106^). Our results suggest that several features of spindles may be required for accurate assays and prediction of intelligence. For example, spindle rate may reflect cortical replay and consolidation and frequency may indicate network maturity.

Our study draws from a database of primarily hospital-based recordings, which biases the subjects included towards those that pursued medical evaluation for an unusual event. Our approach did allow detailed clinical chart review to confirm normal neurodevelopment and the absence of neuroactive medications, or neurological or psychiatric diagnoses expected to impact cortical function, which may improve capture of these concerns compared to subjective reporting in population studies. We also note that the findings reported here are consistent with several smaller population-based studies ^62, 64^. While our study included 19-channel EEG montages, which improves upon the spatial sampling of EEGs used in typical polysomnograms, our EEG durations primarily included only short naps. We therefore could not evaluate the dynamics of spindles over the course of several N2 epochs, as would be expected over a full night of sleep. We note that while subtle changes in spindle rate may be present over the course of a full night ^2^, prior work has also found that naps can reliably estimate overnight spindle rate in adults ^107^. Further, we evaluated spindle dynamics over age using a cross-sectional study design. Validation of the dynamics observed here within individuals will require a long-term longitudinal study. Finally, we evaluated spindles from term infants onward and the developmental trajectory of spindles in premature neonates remains poorly characterized. Immature spindle bursts, also called “delta brushes” or delta-beta complexes, can be first observed in premature neonates at 28 weeks postmenstrual age in the form of coupled slow and fast oscillations triggered by sensory input ^108, 109, 110^. Delta brushes are generated by the same thalamocortical circuits as sleep spindles, can be similarly triggered by cortical sensory events, and serve the same roles in consolidating learned experiences ^110, 111, 112, 113^. “Spindle bursts” in premature infants could therefore provide additional insight and an even earlier biomarker of cognitive function, brain maturation, and thalamocortical circuit integrity.

Sleep spindles are a unique neuronal rhythm that reflect and support cognitive function across the lifespan. Spindles are known to follow a non-linear trajectory from late childhood through late adulthood. This work fills in the critical developmental gap of spindle ontogeny and maturation from birth through adolescence. Our findings reveal that neonatal spindle frequency and topology provide early, transient views of the adolescent form of these genetically influenced traits. In contrast, we show that spindle rate follows a maturational trajectory that closely parallels cortical development. These data expand our current knowledge of normal physiological brain development and provide a large normative database to detect deviations in sleep spindles to aid discovery, biomarker development, and diagnosis in cognitive development and pediatric neurodevelopmental disorders.

## Competing Interests

The authors report no conflicts of interest.

## Acknowledgments

This work was supported by NINDS R01NS115868

## Notes

### Competing Interest Statement

The authors have declared no competing interest.

